# Polycomb proteins translate histone methylation to chromatin folding

**DOI:** 10.1101/2022.05.13.491808

**Authors:** Ludvig Lizana, Negar Nahali, Yuri B. Schwartz

## Abstract

Epigenetic repression often involves covalent histone modifications. Yet, how the presence of a histone mark translates into changes in chromatin structure that ultimately benefit the repression is largely unclear. Polycomb group proteins comprise a family of evolutionarily conserved epigenetic repressors. They act as multi-subunit complexes one of which trimethylates histone H3 at Lysine 27 (H3K27). Here we describe a novel Monte Carlo – Molecular Dynamics simulation framework, which we employed to discover that stochastic interaction of the Polycomb Repressive Complex 1 (PRC1) with tri-methylated H3K27 is sufficient to fold the methylated chromatin. Unexpectedly, such chromatin folding leads to spatial clustering of the DNA elements bound by PRC1.

---

Epigenetic repression often involves covalent histone modifications. In many instances, we know which proteins install the modifications and specifically recognize them. Yet, how the presence and recognition of a histone modification translate into changes in chromatin structure that benefit the repression is largely unclear. Epigenetic repression of developmental genes by Polycomb system is essential for all multicellular animals (Piunti and Shilatifard, 2021; Schuettengruber *et al*., 2017). It involves tri-methylation of histone H3 at Lysine 27 (H3K27) by Polycomb Repressive Complex 2 (PRC2) (Czermin *et al*., 2002; Müller *et al*., 2002) and binding of the modified histone by Polycomb Repressive Complex 1 (PRC1) (Fischle *et al*., 2003). In fruit flies *Drosophila melanogaster*, the only organism where this was directly tested, H3K27 methylation is required for the repression (Pengelly *et al*., 2013). It appears to act as the molecular mark assuring that both copies of a target gene remain repressed after the DNA replication (Coleman and Struhl, 2017; Laprell *et al*., 2017).

Mechanisms by which Polycomb complexes repress transcription are not well understood but seem to involve modifications of the chromatin structure. The chromatin of genes repressed by Polycomb complexes (hereafter Polycomb-repressed genes) is folded in an unusual way. It is more compact compared to chromatin of regular inactive and transcriptionally active genes and displays higher degree of intermixing (Boettiger *et al*., 2016). This chromatin structure requires PRC1 and was suggested to involve self-interactions of one of its subunits (Boettiger *et al*., 2016). Whether tri-methylation of H3K27 is implicated in the chromatin folding has not been investigated.

*Drosophila* genes regulated by the Polycomb system contain specialized Polycomb Response DNA Elements (PREs), which serve as high-affinity binding sites for PRC1 and PRC2 (Kassis and Brown, 2013). Polycomb-repressed genes are embedded in broad chromatin domains enriched in tri-methylated H3K27 (Kahn *et al*., 2006; Papp and Müller, 2006; Schwartz *et al*., 2006). Despite this, PREs remain the only sites where PRC1 is stably bound. Together with the observation that PRC1 continues to bind PREs in cells deprived of PRC2 (Kahn *et al*., 2016), this discounts the hypothesis that methylation of H3K27 is used to mark genes for PRC1 recruitment. If tri-methylated H3K27 does not serve to recruit PRC1 to genes, what is this epigenetic mark good for?

It is tempting to hypothesize that the tri-methylation of H3K27 is part of the chromatin folding mechanism and thereby epigenetically marks Polycomb-repressed genes for folding. Testing this hypothesis experimentally is challenging for two main reasons. First, there are many proteins involved, which makes the *in vitro* reconstitution of a Polycomb-repressed gene prohibitively difficult. Second, complex biochemical interactions between PRC1 and PRC2 (Blackledge and Klose, 2021; Kang *et al*., 2015) make *in vivo* genetic knock-out experiments hard to interpret. To circumvent this problem, we took a computational approach and developed a novel Monte Carlo – Molecular Dynamics (MC-MD) simulation framework. The framework uses Large-scale Atomic/Molecular Massively Parallel Simulator (LAMMPS) (Plimpton, 1995) to model the chromatin motion explicitly but treats the binding of PRC1 to PREs probabilistically. This approach has three advantages over explicit simulation of the entire system. First, it requires no assumptions regarding physical properties of PRC1 in the nucleoplasm. Second, it does not rely on the prior knowledge of PRC1 affinity to PREs, which, so far, has not been directly measured. Third, there is a considerable computational gain for not tracking PRC1 complexes unbound to chromatin.

To build the model, we represent the chromatin as a semi-flexible self-avoiding polymer fiber with repeating units (monomers) of 10-nm (nucleosome) size. The polymer contains 360 monomers of four possible types (Fig. 1A): nucleosomes containing H3K27me3 (methylated nucleosomes, green), nucleosomes lacking H3K27me3 (unmethylated nucleosomes, red), PRC1-bound PREs (orange), and unbound PREs (light grey). A Polycomb-repressed locus is located in the center of the polymer and is represented as a stretch of methylated nucleosomes with four embedded PREs (see Supplementary Materials and Methods for additional details).

**FIG. 1.**
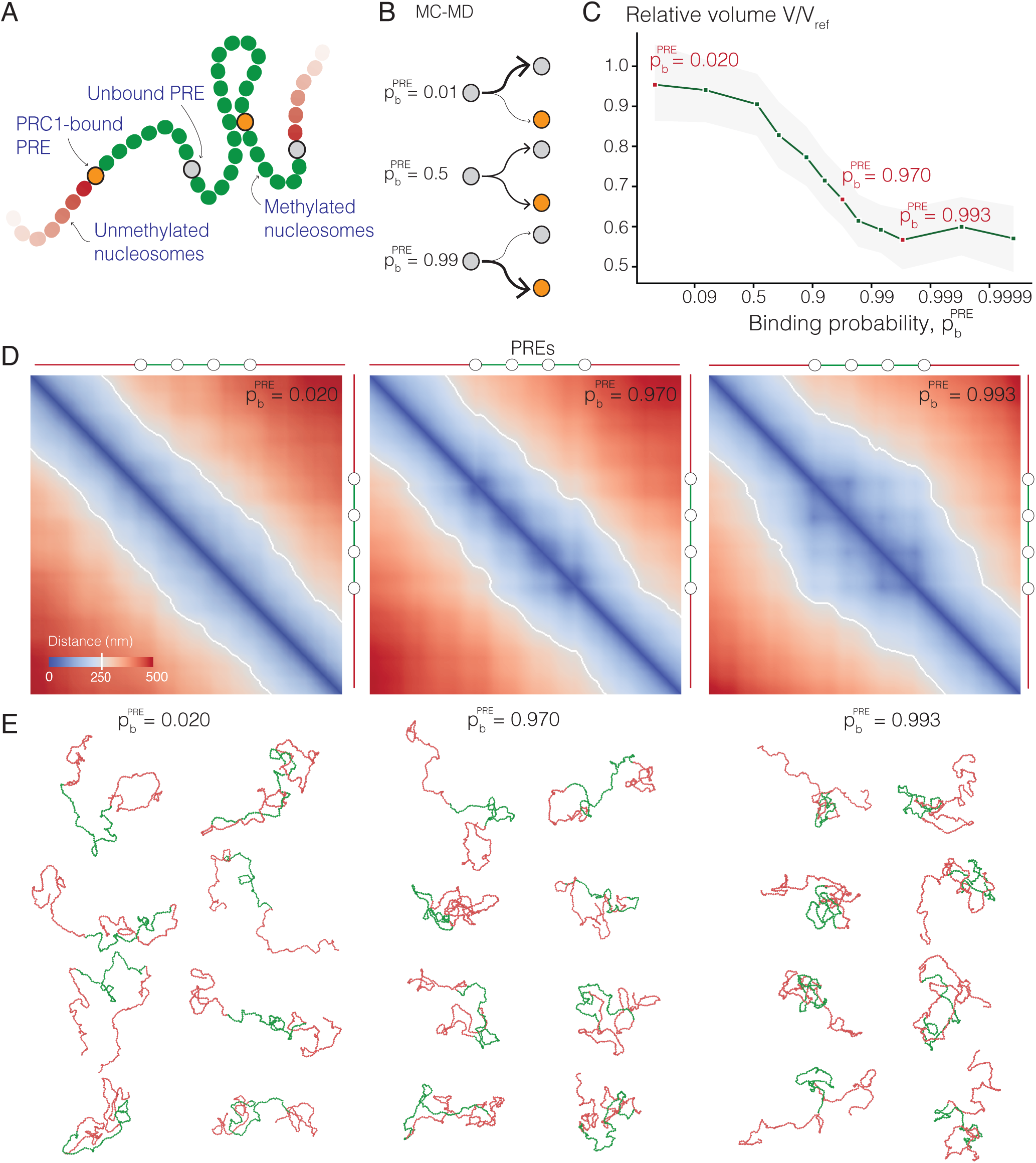
Interactions between PRE-anchored PRC1 and tri-methylated H3K27 fold the chromatin. (A) Polymer model. We model chromatin as a semi-flexible polymer with individual units corresponding to nucleosomes. Nucleosomes come in four types, the most important being PREs that may be PRC1-bound (orange) or free (grey). If bound, PREs have an affinity to nucleosomes tri-methylated at H3K27 (green). Neither PRC1-bound nor free PREs adhere to unmethylated nucleosomes (red). (B) Monte-Carlo – Molecular Dynamics (MC-MD) protocol. Before the polymer MD simulations (using LAMMPS), we probabilistically assign PREs to the PRC1-bound state. The probability of this event (i.e. PRC1 binding probability 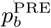) decreases with increasing dissociation constant 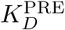. (C) The average simulated volume of the Polycomb-repressed locus at varying 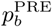 relative to a reference case lacking PREs. The shaded area indicates the 95% confidence interval. (D) Heat-map representation of pairwise distances between monomers for three 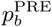 values (indicated in C). At high 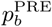, the distance heat-maps display the off-diagonal spots suggesting that PREs come in closer spatial proximity more often than other monomers located at similar linear distances. PRE positions are indicated next to the rightmost panel. (E) Snapshots of representative simulated polymer 3D configurations.

In the first modelling step, we take 150 polymer fibers, organized as described above but with all PREs unbound by PRC1, and simulate their movements with LAMMPS until they reach thermodynamic equilibrium. We then populate the PREs of each equilibrated polymer with PRC1 complexes based on the binding probability 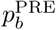 (Fig. 1B). To this effect, we draw uniform random numbers *r* ∈ (0, 1) and designate each PRE as PRC1-bound if 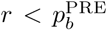. This is followed by a round of LAMMPS simulations that capture events during characteristic residence time (100 sec) of PRC1 on chromatin (Ficz *et al*., 2005). In this simulation round, PRC1-bound PREs are attracted to methylated nucleosomes. If a PRC1-bound PRE and a methylated nucleosome come close, chromatin loops may form. We model the attraction between PRC1-bound PREs and H3K27me3 using the Lennard-Jones potential. The potential was calibrated to recapitulate the binding affinity between PRC1 and H3K27me3. Two independent approaches were used for this purpose. First, we used methods of statistical mechanics to derive the equation linking the Lennard-Jones potential to dissociation constant. Second, we ran a series of designated many-particle simulations using a range of Lennard-Jones potentials to select the one at which the fraction of bound PRC1-H3K27me3 particles matched the dissociation constant 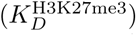 measured experimentally (see Supplementary Materials for details). The two approaches yielded similar results and the average of the two values was used for further analyses.

To understand whether contacts between PRC1-bound PREs and methylated nucleosomes may lead to excessive chromatin folding of the Polycomb-repressed locus, we run the simulations at different 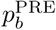 and calculated the volume of the central part of the polymer 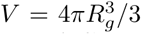 using Radii of Gyration (*R*_*g*_). The Radius of Gyration is defined as the root mean square distance from the polymer’s center of mass to all monomers. Plotting *V* in relation to the average volume of a polymer lacking PREs 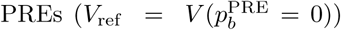 reveals that the volume of the Polycomb-repressed locus shrinks with increasing 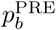 eventually dropping to approximately half of the initial unfolded state (Fig. 1C). The extent of shrinking agrees well with super-resolution microscopy measurements, which indicate that the chromatin of the Polycomb-repressed genes has 40-60% lower volume compared to that of the transcriptionally inactive genes not subjected to the repression (Boettiger *et al*., 2016). Consistently, the heat-map representations of the average pairwise distances between all monomers indicate that those within the Polycomb-repressed locus become shorter as 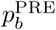 grows (Fig. 1D). The snapshots of individual polymer fibers confirm the trend but also indicate that there is considerable variation between individual fibers even at high 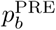 (Fig. 1E).

To summarize, when PREs are bound by PRC1 most of the time, the interactions of the PRE-anchored PRC1 with tri-methylated H3K27 fold the surrounding chromatin. Within the range of the binding probabilities 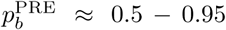, the degree of chromatin folding changes significantly in response to small 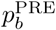 fluctuations. How does this range relate to the probability of PRC1 binding to PREs inside the *Drosophila* cell nucleus? The probability depends on the PRC1 concentration (*c*_PRC1_) and the strength of PRE-PRC1 binding reflected by the dissociation constant 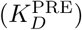. Assuming that the number of PRC1 complexes in the nucleus is substantially higher than the number of PREs (for experimental support of the assumption see: (Bonnet *et al*., 2019; Kahn *et al*., 2016; Schwartz *et al*., 2006, 2010; Steffen *et al*., 2013)), we can view the binding of PRC1 to PREs as pseudo first-order reaction [PRE] + [PRC1] ↔ [PRE : PRC1] and express the binding probability as (see supp. text for derivation):

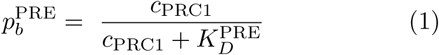

Two independent studies estimated *c*_PRC1_ in *Drosophila* embryonic nuclei as 0.1 - 0.3 *μ*M (Bonnet *et al*., 2019; Steffen *et al*., 2013). Using the average value of *c*_PRC1_ = 0.2 *μ*M in Eq. (1), we see that the binding probability 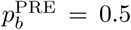 requires PREs to bind PRC1 with 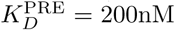, while the 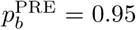 corresponds to 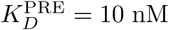. Put another way, compared to the interaction with H3K27me3 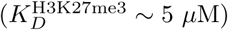 (Fischle *et al*., 2003), PRC1 ought to bind PREs 25 to 500 times stronger. We are not aware of an experimental method to measure 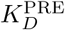 directly. Nevertheless, the difference in the PRC1 binding at PREs and the flanking methylated chromatin measured by Chromatin Immunoprecipitation (ChIP) suggests that such stronger binding of PRC1 to PREs is feasible (Kahn *et al*., 2016). Overall, our calculations and modelling argue that, in *Drosophila* nuclei, PREs are likely to bind PRC1 most of the time and this enables the folding of the surrounding chromatin via interactions of the PRE-anchored PRC1 with tri-methylated H3K27.

The results of our simulations support the hypothesis that the tri-methylation of H3K27 labels Polycomb-repressed genes for chromatin folding. However, to be relevant for the epigenetic transmission of the repressed state, the methylation-driven folding needs to be passed on to daughter copies of a Polycomb-repressed gene following DNA replication. During replication, parental H3 molecules with their posttranslational modifications are randomly partitioned between the two replicating chromatids (Petryk *et al*., 2018; Yu *et al*., 2018). The remainder is supplied via replication-coupled synthesis of unmodified histones. Therefore, immediately after the replication, the density of the H3K27me3-containing nucleosomes drops two-fold and is gradually restored in time for the next replication cycle. Would such dilution of the methylated nucleosomes be compatible with methylation-dependent chromatin folding? To address this question, we repeated MC-MD simulations for three different 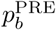 that support chromatin folding. However, this time, we randomly replaced 50% and 25% of H3K27 tri-methylated nucleosomes with unmethylated ones to mimic the situations immediately after the replication and half-way through the re-acquisition of the fully methylated state. As illustrated by Fig. 2, when most PREs are occupied by PRC1 (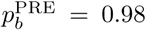or 0.997), even two-fold dilution of methylated nucleosomes has no effect on chromatin folding. When PRC1 binds PREs less often, 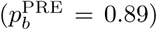, 50% dilution leads to visible increase in the median relative volume of the Polycomb-repressed locus. However, the increase is not significantly different compared to median relative volume of the fully methylated locus. Importantly, the chromatin folding is restored when the density of methylated nucleosomes reaches 75%. To summarise, our simulations argue that tri-methylation of H3K27 is capable to mark Polycomb-repressed genes for epigenetic inheritance of the chromatin folding.

**FIG. 2.**
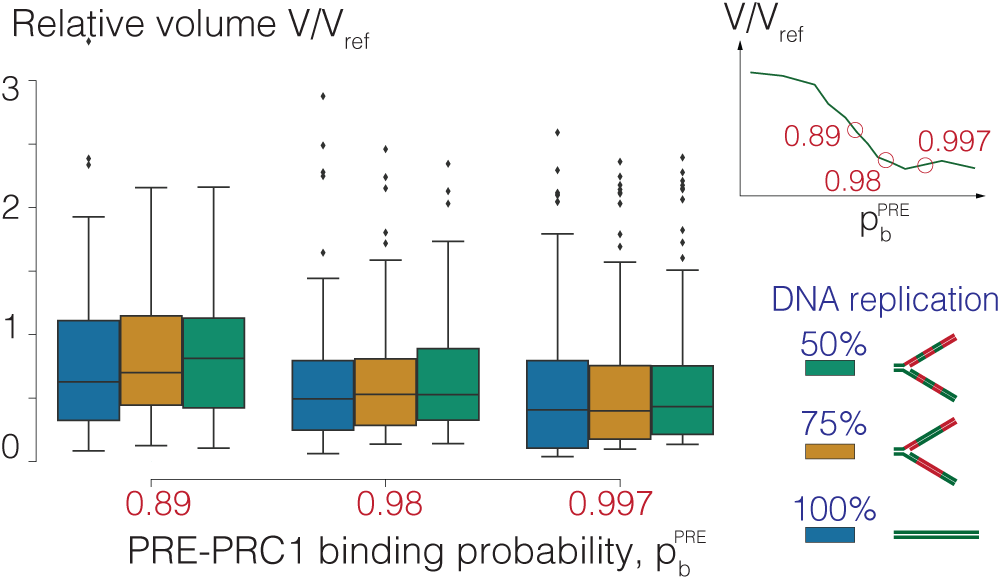
Chromatin folding can be passed to daughter copies of Polycomb-repressed genes after the DNA replication. The boxplots show distributions of relative volumes of Polycomb-repressed loci within 150 polymers simulated at three different probabilities 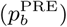. The boxplots indicate the median and span the inter-quartile range. The whiskers show the lowest and the highest values excluding outliers, which are defined as values outside 1.5 inter-quartile range. For each 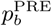, indicated below and on the schematic in the upper right corner, we randomly replaced a fraction of the H3K27me3 nucleosomes with unmethylated ones: 50% methylated nucleosomes left (green) and 75% methylated nucleosomes left (orange). The blue boxplot shows the variation without nucleosome replacement. Note that removing a significant fraction of methylated nucleosomes does not prevent the folding. The relative volumes are scaled to a reference case lacking PREs (*V/V*_ref_).

Besides the overall folding of the Polycomb-repressed locus, the distance heat-maps reveal off-diagonal spots (Fig. 1D). Best seen on the maps for 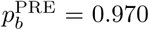 and 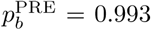, these spots suggest that, when the chromatin of the repressed locus folds, PREs end up close to each other more frequently than other monomers located at comparable linear distances. Strikingly, similar spots were noted in the contact maps of some of the Polycomb-repressed genes assayed by Chromosome Conformation Capture (Hi-C) (Eagen *et al*., 2017; Ogiyama *et al*., 2018). Interpreted as bases of chromatin loops formed by PRE clusters, these spots were hypothesized to arise from the protein-protein interactions between PRE-anchored PRC1 complexes (Eagen *et al*., 2017; Ogiyama *et al*., 2018). Although consistent with the biochemical properties of PRC1 subunits *in vitro*, this explanation cannot apply to our model. Our model does not explicitly simulate PRC1 and, therefore, provides no possibility for direct PRE-PRE interactions. Instead, the PRE clustering appears to have a geometric/probabilistic explanation, which we present below.

Let us first consider a point (e.g. PRC1-bound PRE) on a line (Fig. 3A) that may touch surrounding line segments and form a loop. Depending on the segment type, which could be either “sticky” (e.g. H3K27 trimethylated chromatin) or “non-sticky” (e.g. unmethylated chromatin), the loop will be long-lived or short-lived. The likelihood to touch a specific segment point depends on its distance from the anchor point. The exact distance dependence is not critical for our argument. Therefore, for simplicity, we assume that the loop lengths follow a Gaussian distribution and say that only long-lived loops are stable enough to be detected. We then use a Monte Carlo simulation to model the behaviour of two anchor points in three basic configurations (for additional details see Supplementary text). In the first configuration, the “sticky” segments surround both anchor points. As a result, those may form stable loops to the left and right with the same probability. As illustrated by Fig. 3B, the histograms of resulting loop-lengths are symmetric as is the distribution of the absolute distances Δ between the two anchor points. Consistently, the average distance between the anchor points 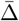 equals 2, which is identical to their linear separation because the stable loops are equally likely to form in any direction. In the second configuration, the leftmost segment is “non-sticky”, while other segments remain the same as in the first configuration. As in this arrangement no long-lived loops form towards the leftmost segment (Fig. 3B, middle row), the histogram of the loop-lengths becomes asymmetric and the average distance between the anchor points shortens to 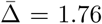. In other words, compared produce the clustering of *inv-en* PREs in our MC-MD to the first configuration, the anchor points become statistically closer. Finally, we consider the configuration where only the segment between the two anchor points is “sticky” (Fig. 3B, bottom row). In this case, stable loops may form only in the direction towards the other anchor point. As a result, the average distance shortens even more to 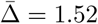. The latter argues that the propensity to cluster is the strongest for PREs located close to the edges of an H3K27me3 domain.

**FIG. 3.**
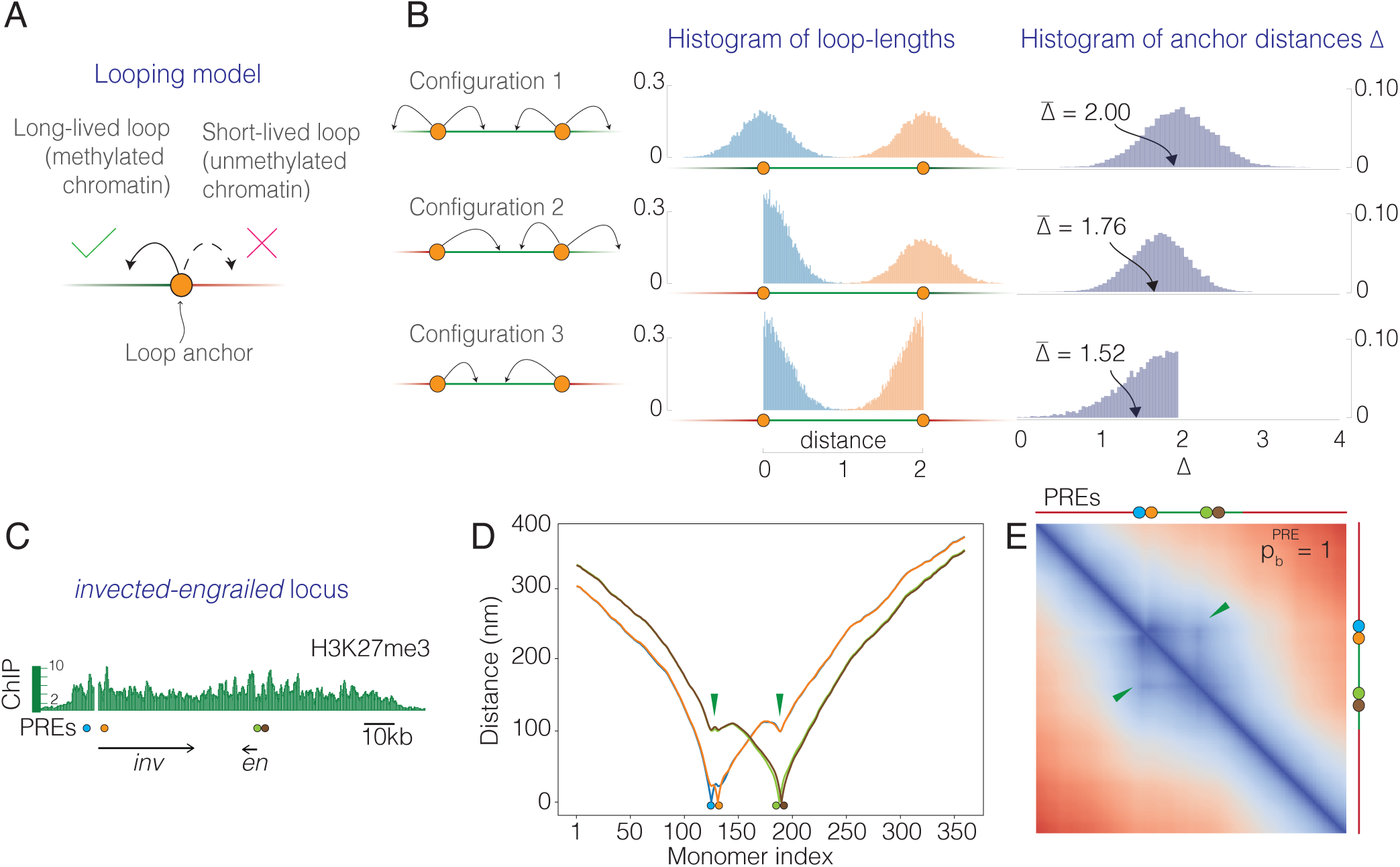
PRE clusters emerge during chromatin folding. (A) Simple looping model. The model considers anchor points (orange circles) on a line that may touch surrounding line segments and form loops. Depending on the segment type, which could be either “sticky” (green) or “non-sticky” (red), the loops will be long-lived or short-lived. The model postulates that only long-lived loops are stable enough to be detected. (B) Three basic configurations of anchor points and the two chromatin types. Arrows indicate the directions in which the anchor points may form the long-lived loops. Shown to the right are the corresponding histograms of the simulated loop-lengths and anchor distances. The average anchor distance 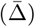 depends on the relative arrangement of anchor points and the “sticky” chromatin segments. (C) The map of the *invected-engrailed (inv-en)* locus repressed by Polycomb system. The distribution of H3K27me3 ChIP signal (13) across the locus indicates the extent of methylated chromatin relative to PREs (blue, orange, green and brown circles) and *inv* and en transcription units (arrows). (D) Color-coded curves show the average 3D distances between individual PREs (marked with circles) and the rest of the monomers (indexed from the left to the right edge of the polymer). The 3D distances between a PRE and other monomers increase with their separation along the polymer. However, green arrows point to valleys, which indicate that distances become smaller, compared with those to the preceding or the subsequent monomers, if the other monomer is also a PRE. (E) Heat-map representation of the pairwise distances between the monomers within the simulated *inv-en* locus. The arrows point to off-diagonal spots formed by PREs.

Interestingly, the *invected-engrailed (inv-en)* locus whose four PREs appear clustered in Hi-C experiments, has this kind of configuration (Eagen *et al*., 2017; Ogiyama *et al*., 2018). We therefore attempted to resimulation framework. To this effect, we repositioned PRE monomers within the simulated polymer such that the relative distances between them and the edges of the Polycomb-repressed gene were the same as in the *inv-en* locus (Fig. 3C) and performed the MC-MD simulations in the same way as described above. Measurements of the distances between each PRE and other monomers (Fig. 3D) or the heat-map of the pairwise distances between monomers (Fig. 3E) indicate that the simulated PREs cluster. Overall, we conclude that looping interactions between PRE-anchored PRC1 and trimethylated H3K27 automatically increase the likelihood that PREs are found in closer proximity compared to other monomers located at similar linear distances.

Two main conclusions follow from the observations presented here. First, in the milieu of *Drosophila* nuclei, the stochastic interactions of the PRE-anchored PRC1 with tri-methylated H3K27 are sufficient to fold the methylated chromatin. This effectively translates the epigenetic marking of the Polycomb-repressed genes into chromatin folding. Conceivably, such folding competes with processes required for transcriptional activity, for example, chromatin looping required for enhancer-promoter interactions. Since the extent of the folding depends on the affinities of individual PREs to PRC1 and their relative arrangement, changes of either or both will allow evolutionary selection of combinations tailored for regulation of specific genes. Second, the chromatin folding by PRE-anchored PRC1 leads to spatial clustering of the DNA elements to which PRC1 is bound. Remarkably, the clustering does not require specific protein-protein interactions. It emerges during chromatin folding and depends on relative positions of PREs inside the Polycomb-repressed genes. Such clustering may be reinforced by interactions between PRC1 complexes (Gambetta and Müller, 2014; Kim *et al*., 2002, 2005). We note that probabilistic clustering does not depend on the molecular nature of the anchor points or chromatin “stickiness”. For example, enhancer elements bound by proteins that interact with elongating RNA polymerase complexes may perceive intensively transcribed genes as stretches of “sticky” chromatin. This, in turn, will lead to enhancer clustering. Decades of prior experimental studies gave us the opportunity to calibrate our Monte Carlo – Molecular Dynamics simulations such that they yielded a reasonably realistic representation of the *Drosophila* Polycomb-repressed gene. While very few other epigenetic systems are investigated to comparable detail, as our understanding of these systems grows, the MC-MD approach presented here will likely be useful to explore them.

## Acknowledgments

We are grateful to Dr. Jan Larsson (Umeå University) for critical reading of the manuscript. This research was conducted using the resources of High Performance Computing Center North (HPC2N). We acknowledge financial support from the Swedish Research Council (grant numbers: 2021-04435 (YBS), 2017-03848 (LL), and 2021-04080 (LL)), Cancerfonden 19 0003 Pj (YBS), and Knut and Alice Wallenbergs Stiftelse 2014.0018 (to EpiCoN, YBS co-PI).

