## Supplementary figures and methods for "Polycomb proteins translate histone methylation to chromatin folding"

Ludvig Lizana*

*Integrated Science Lab, Department of Physics, Umeå University, SE-901 87 Umeå, Sweden*

Negar Nahali

*Integrated Science Lab, Department of Physics, Umeå University, SE-901 87 Umeå, Sweden and Centre for bioinformatics, Department of Informatics, University of Oslo, NO-0373 Oslo, Norway*

Yuri B. Schwartz†

*Department of Molecular Biology, Umeå University, SE-901 87 Umeå, Sweden*

(Dated: May 13, 2022)

### I. MATERIALS AND METHODS

#### A. Polymer model

We describe chromatin using the standard biopolymer model from (Kremer and Grest, 1990). The model builds on three types of interactions associated with stretching, bending, and excluded volume preventing two monomer from occupying the same space. We review each of these contributions below.

*1. Stretching – finitely extensible spring* We model stretching between two neighboring monomers with the non-linear Warner spring (Warner Jr, 1972), also known as the FENE potential:

$$V_{\mathrm{FENE}}(r) = \begin{cases} -\frac{1}{2}K_s\left(\frac{R_0}{\sigma}\right)^2 \ln\left[1-\left(\frac{r}{R_0}\right)^2\right] & r \le R_0 \\ \infty & \text{otherwise,} \end{cases} \quad (1)$$

Here, $r$ denotes the absolute distance between two monomer centers, $K_s$ is the potential's strength, $R_0$ is the spring's maximum extension, and $\sigma$ is the nucleosome diameter ($\sigma = 10$ nm). We use $R_0 = 1.5\sigma$ (Annunziatella *et al.*, 2016) and $K_s = 30\,k_BT$, where $k_B$ is Boltzmann's constant and $T$ is absolute temperature, which gives the average bond length $\approx 0.97\sigma$.

*2. Bending – polymer stiffness.* To describe the stiffness, we use a bending potential that is proportional to the cosine of the angle $\theta$ between neighboring bonds (i.e. the angle between next-nearest-neigbor monomers)

$$V_{\mathrm{bend}}(\theta) = K_\theta\,(1-\cos(\theta)) \quad (2)$$

We set the bending constant to $K_\theta = 3k_BT$. This corresponds to the persistence length $l_p = K_\theta\sigma/(k_BT) = 3\sigma = 30$ nm (Langowski, 2006). In our simulations, we have on average $l_p \approx 2.7\sigma$.

*3. Excluded volume – no monomer-monomer overlap.* We model excluded volume interactions between two monomers from a truncated version of the Lennard-Jones potential

$$V_{\mathrm{LJ}}(r) = 4\epsilon\left[\left(\frac{\sigma}{r}\right)^{12} - \left(\frac{\sigma}{r}\right)^{6}\right], \quad (3)$$

where, $\epsilon \ge 0$ is the potential's strength and, as before, $\sigma$ is the monomer diameter and $r$ the absolute distance. However, as $V_{\mathrm{LJ}}$ includes both repulsion and attraction, we turn it into an excluded volume interaction, $V_{\mathrm{EV}}$, by cutting the potential at the minimum ($r = 2^{1/6}\sigma$) and shifting it upwards by $\epsilon$. For distances beyond the cut, the interaction energy is zero (Fig. 1, yellow curve). In other words,

$$V_{\mathrm{EV}}(r) = \begin{cases} V_{\mathrm{LJ}}(r) + \epsilon & r \le 2^{1/6}\sigma \\ 0 & \text{otherwise.} \end{cases} \quad (4)$$

We put $\epsilon = 1\,k_BT$ to get a strong enough repulsion.

*Monomer-monomer attraction.* In addition to the standard polymer properties – stretching, bending, and

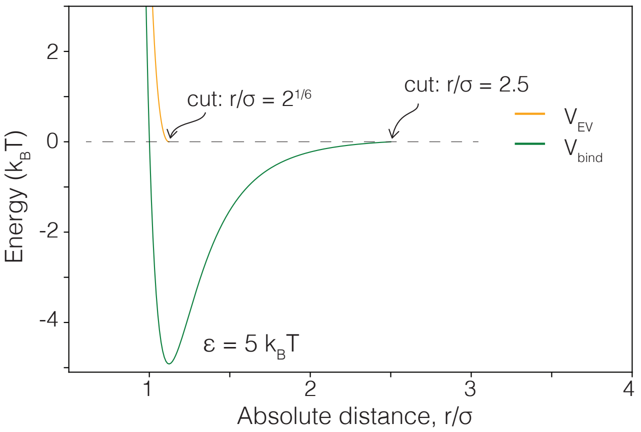


FIG. 1 **Attraction and repulsion potentials in the polymer model.** The curves show the binding energy $V_{\mathrm{bind}}(r)$ (green) and the excluded-volume repulsion energy $V_{\mathrm{EV}}(r)$ (orange) when $\epsilon = 5k_BT$.

---

*Electronic address:
†Electronic address:

excluded volume interactions – some monomers also attract each other. Similar to (Annunziatella *et al.*, 2016), we model such attraction using the Lennard-Jones potential (Eq. 3) and assume that the interactions do not extend beyond some interaction distance $r = r_{\rm int}$, after which the binding potential is zero. In summary, we model monomer attractions with

$$V_{\rm bind}(r) = \begin{cases} V_{\rm LJ}(r) - V_{\rm LJ}(r_{\rm int}), & r \le r_{\rm int} \\ 0, & \text{otherwise} \end{cases}, \quad (5)$$

where $r_{\rm int} = 2.5\sigma$. We show $V_{\rm EV}(r)$ and $V_{\rm bind}(r)$ in Fig. 1 for easy comparison.

*Equation of motion.* Just as in (Kremer and Grest, 1990), we assume that each monomer undergo Brownian motion and subject to a force $\mathbf{F}$ associated with the interaction potentials specified above. Denoting $\mathbf{r}_i$ as the $i$th monomer's 3D coordinate and $V(\mathbf{r}_i)$ as the total interaction potential (sum of Eqs. (1)-(5)), the force is $\mathbf{F}_i = -\nabla V(\mathbf{r}_i)$. We use the Langevin equation for the monomers' equation of motion.

$$m\frac{d^2\mathbf{r}_i(t)}{dt^2} = -\Gamma\frac{d\mathbf{r}_i(t)}{dt} + \mathbf{F}_i + \mathbf{W}_i(t) \quad (6)$$

where $\Gamma$ is the damping constant coupling the monomers to the surrounding fluid and $\mathbf{W}_i(t)$ is a white-noise force term. The noise's amplitude is related to $\Gamma$ via the diffusion diffusion constant $D$, using the Einstein relation $D = k_B T/\Gamma$. We simulate Eq. (6) using LAMMPS (see Sec. I.E).

### B. Polymer configuration and monomer types

We simulate a polymer consisting of 360 monomers. As each monomer represents a nucleosome, the whole polymer corresponds to approximately a 63 kb chromatin segment.The polymer is divided into three sections (see Fig. 2). The middle third contains H3K27me3 nucleosomes (green) with a few interspersed PREs (grey or orange). The two other thirds hold only unmethylated monomers (red).

PREs come in two flavors: PRC1-bound (orange) or unbound (grey). If bound, they attract methylated monomers and can form long-lived loops if appearing close enough in 3D. We do not consider attraction between PRC1-bound PREs and unmethylated monomers (red). Before the polymer simulation starts, we assign PRC1s randomly to the PREs. We outline this procedure below in Sec. I.C.

We adjusted the flanking regions' length (and the volume of the simulation box) to calibrate the monomer volume fraction to the nucleosome density in the nucleus. In our simulation, we use 360 monomers, each having size $\sigma = 10$ nm, residing in a total volume that is $(24\sigma)^3$ large. This yields the volume fraction $360/24^3 = 0.026$. This is very close to the nuclear volume fraction that we estimates as follows. First, the volume of *Drosophila*'s cell nucleus is 78 $\mu$m$^3$. It holds approximately 1.65 million nucleosomes (the total DNA length is $2.88 \cdot 10^8$ bp and each nucelsome contains 175 bp). Second, partitioning the nuclear volume into nucleosomes-sized 10 nm subvolumes, each having volume $10^{-6}\mu$m$^3$, there are $78 \cdot 10^6$ of such volumes. Given 1.65 million nucleosomes, the filling fraction is $1.65 \cdot 10^6/78 \cdot 10^6 = 0.021$.

### C. Monte Carlo - Molecular Dynamics Scheme (MC-MD)

Instead of simulating diffusing PRC1 proteins surrounding the polymer searching for target monomers, we use binding probabilities to assign PRC1s randomly to the PREs prior to the polymer simulation. We start by making the assignment, and then introduce interaction potentials and simulate the polymer's motion in LAMMPS. Below, we outline are the main steps in our numerical approach. See also Fig. 2.

1. Set binding probabilities $p_b^{\rm PRE}$, $p_b^{\rm me3}$, and $p_b^{\rm unmeth.}$.

2. Sequentially go through all monomers $i = 1, ..., 360$. For each monomer:

   (a) Draw a random number $r$ uniformly distributed between 0 and 1.

   (b) Populate the monomers with PRC1s based on $r$ and the binding probabilities. We consider two cases:

      i. If the $i$th monomer is a PRE and $r < p_b^{\rm PRE}$, we introduce the binding potential $V_b$ (Eq. (5)) between the PRE and all methylated monomers.

      ii. If the $i$th monomer is a methylated or unmehtylated nucleosome, we leave it unbound. In other words, we assume that $p_b^{\rm me3} = p_b^{\rm unmeth.} = 0$. However, we point out that this choice is not a limitation of our framework and nothing prevents us from considering non-zero binding probabilities and introducing weak interactions potentials.

3. Run polymer simulations. After we assigned PRE with PRC1s, we simulate the polymer in LAMMPS. At regular time intervals, we store the monomers' 3D coordinates that we use to calculate averages. We stop the simulation after 100 sec ($\sim 10^9$ LAMMPS MD steps) when we consider the PRC1s as unbound.

We point out that we calculte averages over uncorrelated polymer snapshots and repeated PRE assignments keeping $p_b^{\rm PRE}$ constant. That is, first we set $p_b^{\rm PRE}$, then we collect data as we cycle through the above items 2-3 several times, and then we calculate averages.

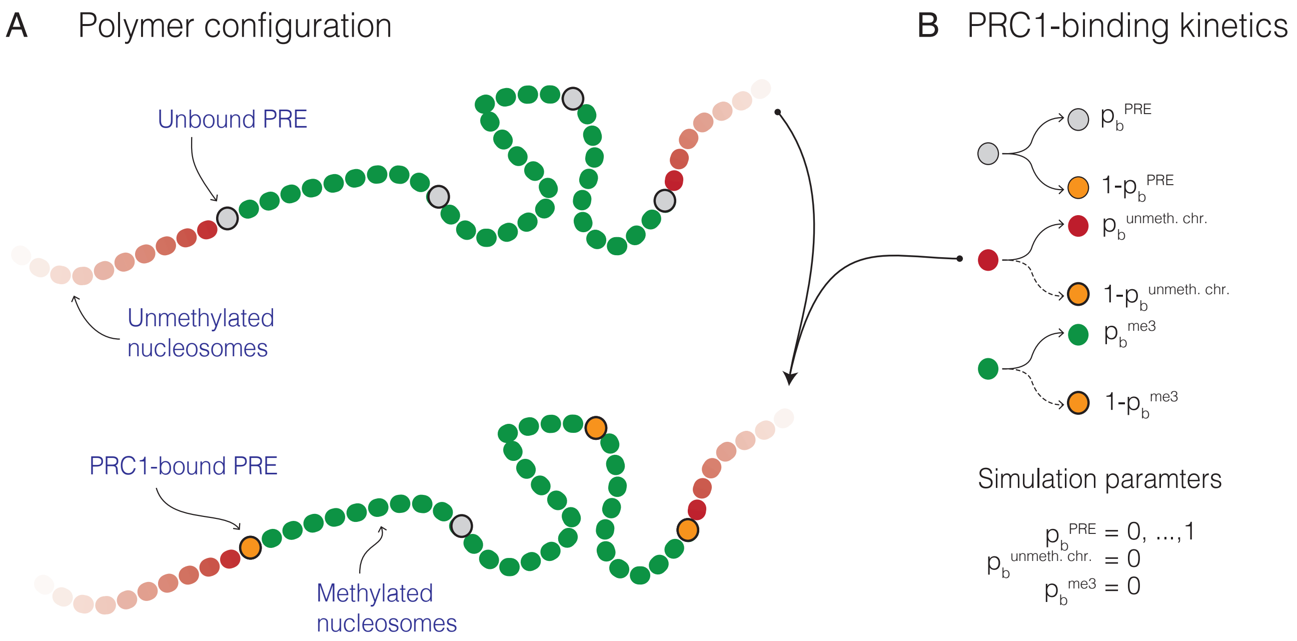


FIG. 2 (A) **Polymer configuration.** The upper polymer contains three monomer types where all PREs are unbound (gray-filled circles). The lower polymer show that some PREs are populated by PRC1 complexes. This is assignment is random and follow the binding scheme in (B) (outlined in Sec. I.C).

#### D. Equipment

We ran all our simulations on the High Performance Computing Center North (HPC2N),[1] a center for Scientific and Parallel Computing. We used $\lesssim 5$ computer nodes each having 128 GB of memory and $2 \times 14$ cores (CPU: Intel Xeon E5-2690v4). Depending on binding strengths, the polymer equilibration took $\sim 3-7$ days. The ensuing simulation where we store the polymer's 3D coordinates, took $\sim 1-3$ days.

#### E. Simulation details: system preparation, equilibration and simulation time

To study the polymer fluctuations, we used the LAMMPS molecular dynamics package. We included interaction potentials specified in Eqs. (1)-(5) and then simulated the polymer under fixed temperature and volume conditions with periodic boundary conditions. Furthermore, we used a Langevin thermostat to treat the surrounding water as an implicit solvent and integrated the monomers' equation of motion (Eq. (6)) using a velocity Verlet algorithm. Below, we outline critical steps in our simulation scheme.

[1] https://www.hpc2n.umu.se

##### 1. Simulation time scales

We integrate Eq. (6) using the velocity Verlet algorithm. Like in (Annunziatella *et al.*, 2016), we set the MD timescale $\tau$ by considering the diffusion constant $D$. Using $D = 5\,(\mu\text{m})^2\text{s}^{-1}$, which is reasonable for a 10 nm macromolecule, gives

$$\tau = \frac{\sigma^2}{D} = \frac{(10\,\text{nm})^2}{5\,\frac{(\mu\text{m})^2}{\text{s}}} = 20\mu\text{s} \qquad (7)$$

In terms of $\tau$, we set damping term and integration step to $\Gamma = 0.5\tau^{-1}$ and $\Delta t = 0.012\tau$.

##### 2. System preparation

Before starting the simulation, we remove all monomer-monomer overlaps. This step ensures that force gradients do not diverge, causing the simulation to crash. We achieve this in three steps. First, we randomly placed the monomers in a giant simulation box, much bigger than we used later on. Second, we push overlapping monomers apart using the soft potential

$$\phi = A\left[1 + \cos\left(\frac{\pi r}{r_c}\right)\right], \quad r < r_c, \quad \phi = 0, \text{ otherwise} \qquad (8)$$

where $r_c = 2.5\sigma$, and where we increased $A$ from 0 to 40 in linear increments during a short MD run (just a few MD steps). In the third step, we removed $\phi$ and made a quick run (about 500 MD steps) to reduce the volume and achieve the desired monomer density (and pressure). After these steps, we have a non-overlapping polymer inside the target volume.

##### 3. Equilibration

After achieving the desired density, we let the polymer relax to an equilibrated state with respect to the interaction potentials we specified in Eqs. (1)-(4)) (excluding monomer-monomer attractions). The equilibrium process took $2 \times 10^6$ MD steps under fixed volume and temperature conditions, and periodic boundaries. The simulation time is long enough so that the polymer's center-of-mass starts to diffuse, indicating that all internal degrees of freedom are equilibrated, such as single monomer fluctuations.

##### 4. Simulation time

After completing the equilibration step, we populate some PREs with PRC1 complexes (outlined in Sec. I.C) and introduce positive attractions accordingly using Eq. (5). Then we simulate the polymer's fluctuations and sample its 3D configuration at regular (non-correlated) time intervals. We stop the simulation after $100 \times 10^6$ MD steps, corresponding to 100 seconds, which is associated with typical PRC1 residence times (Ficz *et al.*, 2005).

#### F. Volume and the Radius of Gyration

To understand by how much the methylated polymer region folds as we change $p_b^{\mathrm{PRE}}$, we calculated the region's voluming $\mathcal{V}$ using the Radius of Gyration $R_G$, where $\mathcal{V} = \frac{4}{3}\pi R_G^3$. For a polymer with $N$ segments, $R_G$ is the average distance between each monomer and the polymers' center point $\mathbf{r}_c = \frac{1}{N}\sum_{k=1}^{N}\mathbf{r}_k$. That is,

$$R_G^2 = \frac{1}{N}\sum_{k=1}^{N}(\mathbf{r}_k - \mathbf{r}_c)^2, \tag{9}$$

As we only calculate $R_G$ for monomers inside the methylated regions, we restrict the monomer indices to $i = 120, ..., 240$.

### II. SUPPLEMENTARY TEXT

#### A. Binding energies and binding constants

The binding constant $K_D$ depends on the binding energy and entropy cost of forming the bond. Therefore, to ensure that our simulations have the proper $K_D$ and associated binding probability $p_b$, we must balance the concentration of binding proteins $c$ and the Lennard-Jones energy scale $\epsilon$ that sets the binding energy. In this section, we outline two ways how to accomplish this balance. One method uses explicit LAMMPS simulations that resemble the PRC1-H3K27me3 peptide-binding system studied in (Fischle *et al.*, 2003). The other is a theoretical estimate based on statistical mechanics arguments valid for large simulation volumes.

These approaches gives two different estimates: $\epsilon/(k_BT) = 3.94$ (numerical) and $\epsilon/(k_BT) = 6.20$ (theoretical). While both approaches have merit, they are not perfect representations of the actual PRC1-H3K27me3 system. For example, in the numerical simulations, one H3K27me3 'particle' may attract more than one PRC1 'particle'. While this is not inconceivable, it has not been observed in experiments. This contrasts with the theoretical estimate that forbids multiple binding but, on the other hand, assumes that all proteins have the same size. Because both approaches have limitations, we use the average Lennard-Jones parameter to fix the PRC1-H3K27me3 interaction. That is

$$\frac{\epsilon}{k_BT} = \frac{3.94 + 6.20}{2} = 5.07 \approx 5.1 \tag{10}$$

Below, we outline each of the two approaches separately.

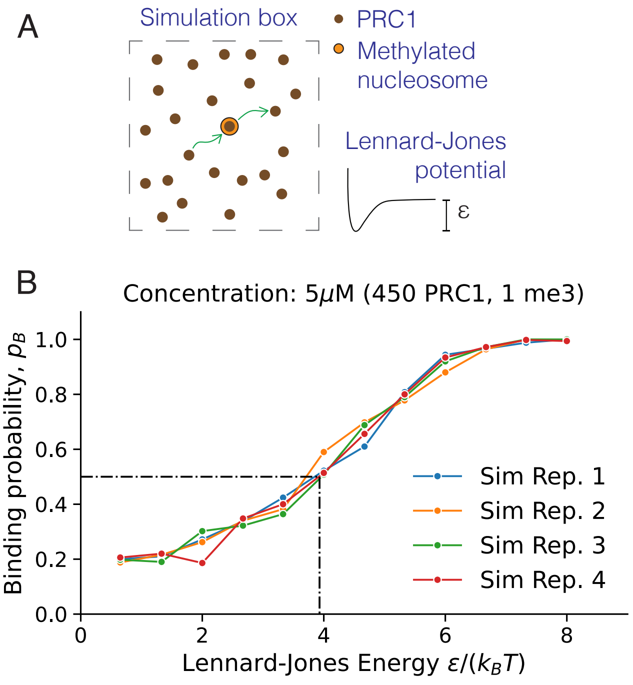


FIG. 3 **Simulated binding probability (bound fraction) for three Lennard-Jones energies $\epsilon$.** (A) Simulation setup where one binding site (orange) is surrounded by diffusing particles (brown). This setup reassembles PRC1 binding to H3K27me3 peptides studied experimentally in (Fischle *et al.*, 2003). (B) Calibration curve. Each line (Rep. 1-4) shows four different simulation results with the same parameters (450 particles, 1 binding site). The dots in each line represent the fraction of times the H3K27me3 binding site is bound by at least one PRC1 particle over 500 uncorrelated time points. As indicated with the horizontal dash-dotted line, the simulation parameters are at $K_D$ conditions when the $p_b = 0.5$. That is, $\epsilon/(k_BT) = 3.94$.

##### 1. Numerical estimate of $K_D$

To find the KD conditions using LAMMPS, we simulated a fixed concentration of diffusing particles and one binding site. Then we varied the Lennard-Jones parameter and tracked the fraction of bound proteins – this is a proxy for the binding probability $p_b$ – over several simulation runs. We are at $K_D$ conditions when $p_b = 0.5$.

Figure 3 shows the calibration data. Each line corresponds to four separate simulations ('replicates') having the same particle concentration $c = 5$ $\mu$M. To get good statistics, we used a larger simulation box than in our polymer simulations and thus more particles to achieve the desired concentration (450 particles and $V = (52\sigma)^3$). The dots in each line represents an average of 500 simulated data points. Having four replicates (Rep. 1-4) amounts to 2,000 data points for each value of $\epsilon$. To get the the numerical estimate for the $K_D = 5$ $\mu$M condition, we draw a horizontal line at $p_b = 0.5$. We find that it intersects with the $p_b$ curves when $\epsilon/(k_BT) = 3.94$.

##### 2. Theoretical estimate of $K_D$

Instead of doing simulations, we may estimate $K_D$ theoretically in terms of $\epsilon$ assuming that the simulation volume is large. We derive this estimate in three steps.

First, we calculate the binding energy $E_b$ in terms of $V_b(\epsilon)$ (Eq. (5)). By definition, $E_b$ is the energy it takes to move a test particle from the bottom of the energy well past the interaction distance. Because $V_b$ is constant beyond this distance (zero in our notation, see Fig. 1), moving the test particle even further is not associated with any extra energy cost. Using Eq. (5), $E_b$ becomes

$$E_b = V_{\text{bind}}(r_{\text{min}}) - V_{\text{bind}}(r_{\text{int}}). \tag{11}$$

Using that $r_{\text{min}} = 2^{1/6}\sigma$ and $r_{\text{int}} = 2.5\sigma$, we find that

$$E_b \approx -0.938\epsilon. \tag{12}$$

In the second step, we consider the binding probability $p_b$. It combines the binding energy $E_b$ and entropy cost from bringing a test particle near the binding site. This cost grows with the volume $\mathcal{V}$. From statistical mechanics (e.g., (Phillips *et al.*, 2012)), we have $p_b = (1 + \exp(E_b/(k_BT))/\mathcal{V})^{-1}$. However, if there is a concentration $c$ of particles, this expression modifies to

$$p_b = \frac{c}{c + c_0 e^{E_b/(k_BT)}}, \tag{13}$$

where $c_0$ is standard state. This state is often set to 1 M. However, in our simulations $c_0 = 1.67 \times 10^{-3}$ M.[2]

Lastly, we relate $E_b$ to $K_D$ via $p_b$. From first-order chemical kinetics, one can show that (see Sec. II.C)

$$p_b = \frac{c}{c + K_D} \tag{14}$$

Comparing expressions (13) and (14), we identify that the binding constant is

$$K_D = c_0 e^{E_b/(k_BT)} \approx c_0 e^{-0.938\epsilon/(k_BT)} \tag{15}$$

where we used Eq. (12) in the last step. Finally, rewriting this equation gives

$$\frac{\epsilon}{k_BT} \approx -\frac{1}{0.938} \ln\left(\frac{K_D}{c_0}\right) \tag{16}$$

This equation allows us to calculate the Lennard-Jones parameter $\epsilon$ associated with a desired $K_D$. For example, if $K_D = 5$ $\mu$M and $\mathcal{V} = (23\sigma)^3$, then $\epsilon = 6.20$ $k_BT$.

### B. Minimal model for chromatin looping

In Fig. 1C (main text), we argue that the methylated chromatin regions fold with increasing binding probability $p_b^{\mathrm{PRE}}$. In addition, we observe that off-diagonal spots appear in the distance maps (Fig 1D), suggesting that PREs come in proximity more often than their flanking methylated nucleosomes do. This seems contradictory because there are no explicit PRE-PRE interactions in the model. However, there is a simple geometric explanation to why these spots emerge. Here we construct a simple mathematical model for chromatin looping showing the underlying principles.

Consider a loop anchor on a line we show in Fig. 3 (main text) that may touch surrounding line segments and form a loop. Depending on the surrounding segment types only some loops are long-lived enough to count as stable. We consider two types representing methylated (green) and unmethylated (red) nucleosomes. If the anchor reaches out to a green part, we consider the loop as stable. But if that part instead is red, we treat it as unstable, and the loop does not form. This scenario mimics stable interactions between PRC1 and H3K27 methylated nucleosomes and transient interactions in between PRC1 and unmethylated chromatin. Below we refer to green and red segments as sticky and non-sticky.

Apart from segment type, the probability to form a stable loop depends on the loop length. However, the exact length dependence is not crucial for this analysis. It is more important how the loop anchors are positioned relative to different types of segments that restricts some looping directions. Therefore, going forward, we assume that the loop lengths $l$ follow a Gaussian distribution

$$g(l) = \frac{\exp\left(-(l - l_0)^2/(2\sigma_l^2)\right)}{\sqrt{2\pi\sigma_l^2}} \tag{17}$$

where $l_0$ denotes the loop anchor's position.

In this analysis, we track the absolute distance $\Delta$ between two loop anchors after looping. We expect that the distance distribution, $p(\Delta)$, changes with the loop anchors' position relative to sticky (methylated) and non-sticky (unmethylated) segments. We study three configurations shown in Fig. 3 (main text).

#### 1. Configuration 1: embedded loop anchors

In this configuration, sticky segments surround both anchor points – like the two embedded PREs in our polymer model. These may form loops to the left and right with the same probability. We put the loop anchors at positions $l_{10} = 0$ and $l_{20} = 2$, so that $\Delta = 2$ whenver there is no loop, and draw random loop lengths from

$$\begin{aligned} g(l_1) &= \frac{\exp\left(-l_1^2/(2\sigma_l^2)\right)}{\sqrt{2\pi\sigma_l^2}}, \\ g(l_2) &= \frac{\exp\left(-(l_2 - 2)^2/(2\sigma_l^2)\right)}{\sqrt{2\pi\sigma_l^2}} \end{aligned} \tag{18}$$

where $\sigma_l = 0.2$. To mimic many looping events, we draw $10^4$ random numbers from these distributions and calculate the absolute distance

$$\Delta = |l_1 - l_2| \tag{19}$$

Figure 3 (main text) shows the normalized loop-length histograms surrounding each anchor point, and the distribution of the absolute distances $\Delta$. We see that $p(\Delta)$ is symmetric with respect to the average $\bar{\Delta} = 2$ ($\bar{\Delta} = \int_0^\infty p(\Delta)d\Delta$). This is identical to the linear anchor-point separation because loops may form in any direction.

#### 2. Configuration 2: one embedded anchor point and one on the edge

Here, we replace the leftmost segment in Configuration 1 with a non-sticky segment (red). This restricts the left anchor point so it can no longer form loops to the left, only to the right (towards the other loop anchor). In our polymer setup associated with Fig. 1 (main text),

[2] The concentration of one 10 nm particle in the simulation volume $\mathcal{V} = (23 \times 10\,\mathrm{nm})^3$ is 0.12$\mu$M. Thus, the concentration of a filled volume with $24^3$ particles is $c_0 = 0.12\,\mu\mathrm{M} \times 24^3 = 1.67$ mM.

this corresponds to the rightmost (or leftmost) PRE pair where one of the PREs rests on the edge of the methylated region and the other one is inside.

In this setup, we cannot use Eq. (17) for the left anchor point as it assumes that the loops are symmetric around $l_0$. Instead, we use a half Gaussian. For the right anchor, we use the same Gaussian as in Configuration 1.

$$\begin{aligned} g_{\frac{1}{2}}(l_1) &= \frac{\exp\left(-l_1^2/(2\sigma_l^2)\right)}{\sqrt{\pi\sigma_l^2/2}}, \quad l_1 > 0 \\ g(l_2) &= \frac{\exp\left(-(l_2-2)^2/(2\sigma_l^2)\right)}{\sqrt{2\pi\sigma_l^2}} \end{aligned} \tag{20}$$

As the loops are restricted from the left flank, there is a shift in $p(\Delta)$. It is no longer symmetric and the average distance shrinks from $\bar{\Delta} = 2$ (Configuration 1) to $\bar{\Delta} = 1.76$. In other words, the loop anchors are statistically closer to each other in Configuration 2 than in Configuration 1.

#### 3. Configuration 3: loop anchors on the edges

Here, we also exclude loops to the far right, considering also the rightmost segment as non-sticky. In our polymer simulations, this is similar to the PREs on the methylated region's edge. For this case, we draw loop lengths from two half-Gaussians

$$\begin{aligned} g_{\frac{1}{2}}(l_1) &= \frac{\exp\left(-l_1^2/(2\sigma_l^2)\right)}{\sqrt{\pi\sigma_l^2/2}}, \quad l_1 > 0 \\ g_{\frac{1}{2}}(l_1) &= \frac{\exp\left(-(l_2-2)^2/(2\sigma_l^2)\right)}{\sqrt{\pi\sigma_l^2/2}}, \quad l_2 < 2 \end{aligned} \tag{21}$$

This setup shifts $\bar{\Delta}$ even more: from $\bar{\Delta} = 2$ (Configuration 1) and $\bar{\Delta} = 1.76$ (Configuration 2), to $\bar{\Delta} = 1.52$ (Configuration 3).

We point out that there is a fourth case where the sticky segments are on the outer flanks. We did not analyze this case because it will, on average, move PREs further apart from each other. We are mainly interested in situations bringing them closer.

### C. Derivation of the PRC1-PRE binding probability $p_b^{\mathrm{PRE}}$

Here we will derive Eq. 1 in the main text. First, assuming first order kinetics, we have

$$\mathrm{PRC1} + \mathrm{PRE} \longleftrightarrow \mathrm{PRE}', \tag{22}$$

where PRE′ denotes PRC1-bound PREs. In equilibrium, we have

$$K_D^{\mathrm{PRE}} = \frac{c_{\mathrm{PRC1}} c_{\mathrm{PRE}}}{c_{\mathrm{PRE}'}} \tag{23}$$

where $c_X$ denotes concentration. To get $p_b^{\mathrm{PRE}}$, we calculate the fraction of bound PREs, that is

$$p_b^{\mathrm{PRE}} = \frac{c_{\mathrm{PRE}'}}{c_{\mathrm{PRE}}^{\mathrm{tot}}} \tag{24}$$

where $c_{\mathrm{PRE}}^{\mathrm{tot}}$ is the total PRE concentration

$$c_{\mathrm{PRE}}^{\mathrm{tot}} = c_{\mathrm{PRE}'} + c_{\mathrm{PRE}} \tag{25}$$

Using this formula in Eq. (24), as well as the definition for $K_D^{\mathrm{PRE}}$ [Eq. (23)], we obtain

$$\begin{aligned} p_b^{\mathrm{PRE}} &= \frac{c_{\mathrm{PRE}'}}{c_{\mathrm{PRE}'} + c_{\mathrm{PRE}}} = \frac{1}{1 + \frac{K_D^{\mathrm{PRE}}}{c_{\mathrm{PRC1}}}} \\ &= \frac{c_{\mathrm{PRC1}}}{c_{\mathrm{PRC1}} + K_D^{\mathrm{PRE}}} \end{aligned} \tag{26}$$

This is Eq. (1) in the main text.
